# Emergence of opposite neurons in a firing-rate model of multisensory integration

**DOI:** 10.1101/814483

**Authors:** Ho Yin Chau, Wen-Hao Zhang, Tai Sing Lee

## Abstract

Opposite neurons, found in macaque dorsal medial superior temporal (MSTd) and ventral intraparietal (VIP) areas, combine visual and vestibular cues of self-motion in opposite ways. A neural circuit recently proposed utilizes opposite neurons to perform causal inference and decide whether the visual and vestibular cues in MSTd and VIP should be integrated or segregated. However, it is unclear how these opposite connections can be formed with biologically realistic learning rules. We propose a network model capable of learning these opposite neurons, using Hebbian and Anti-Hebbian learning rules. The learned neurons are topographically organized and have von Mises-shaped feedforward connections, with tuning properties characteristic of opposite neurons. Our purpose is two-fold: on the one hand, we provide a circuit-level mechanism that explains the properties and formation of opposite neurons; on the other hand, we present a way to extend current theories of multisensory integration to account for appropriate segregation of sensory cues.

## 1 Introduction

Studies have found congruent neurons and opposite neurons in macaque dorsal medial superior temporal area (MSTd) and the ventral intraparietal (VIP) areas [1, 2, 3, 4]. Congruent neurons prefer and combine visual and vestibular cues of the same self-motion (heading), and have been proposed to be the neural basis of multisensory integration in monkeys [3]. On the other hand, opposite neurons, which prefer and combine visual and vestibular cues of opposite self-motion, do not seem to be involved in multisensory integration [3]. While the role of congruent neurons in multisensory integration has been well-studied both theoretically and experimentally [5, 6, 7], the function of opposite neurons in multisensory integration is still unclear.

Recently, however, it has been proposed that opposite neurons are involved in the decision to integrate or segregate visual and vestibular heading cues based on the likelihood that these cues have a common cause [6]. For example, if visual and vestibular heading cues are largely consistent, then it is likely that they have a common cause, and therefore information about heading provided by both cues should be integrated [8, 3]. On the other hand, if a person is wearing a virtual reality headset but sitting still, then visual and vestibular cues would be inconsistent and the brain should segregate cue information [9]. The problem of inferring the cause of the cues is known as *causal inference* [10, 11], and psychological evidence has shown that indeed the brain carries out such an operation [10, 12, 13, 14]. Optimal decision of integration/segregation can be made by computing the likelihood ratio between integration and segregation, also known as the Bayes factor. A recent paper currently under review proposed that opposite neurons could provide a key step in the computation of the Bayes factor [15], suggesting a potential mechanism of causal inference in the brain.

Because of the potential significance of their computational function, it is desirable to understand how these neurons are learned in the brain. Opposite neurons are characterized by their opposite tuning to cues from two different sensory modalities. For example, an opposite could be most strongly tuned to a rightward visual self-motion cue and a leftward vestibular self-motion cue. The simplest formulation of an opposite neuron involves the projections of two oppositely tuned sensory neurons onto the same neuron. While it could explain the tuning properties of opposite neurons, it is unclear how such projections could be learned. Typical Hebbian learning would require correlation between the firing of one sensory neuron and the firing of its opposite counterpart in the other sensory modality. However, this is biologically implausible in the case of self-motion, as we seldom receive visual cues of self-motion that is contradictory with vestibular cues. Even if there is a significant amount of noise such that one input occasionally contradicts the other, which would cause some opposite connections to form via Hebbian learning, the statistical fact that most of the multisensory neurons in the macaque MSTd are either congruent or opposite but not somewhere in between suggests that this is unlikely (see Fig. 1A).

**Figure 1:**
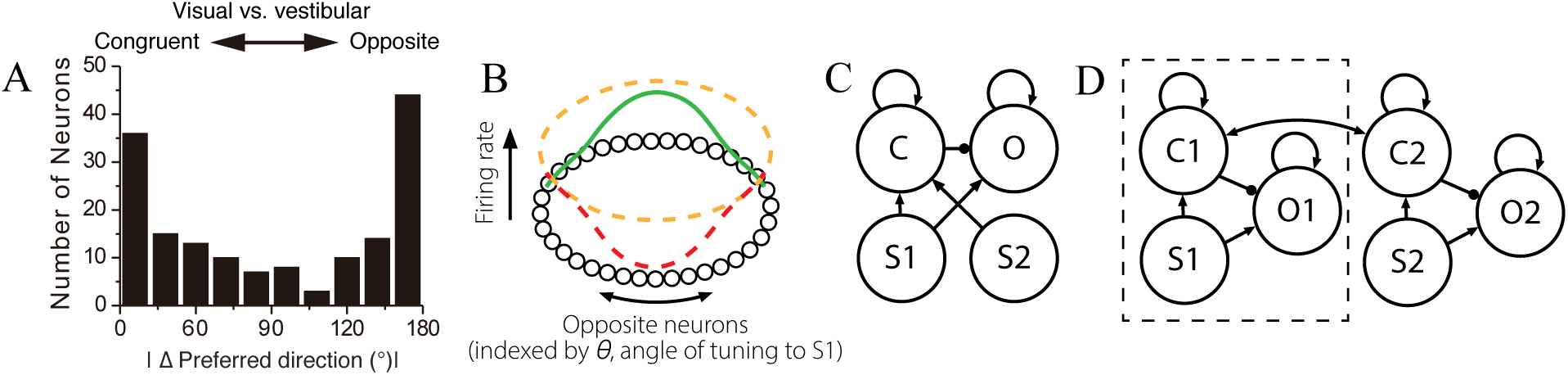
A) Distribution of congruent and opposite neurons in macaque MSTd. Adapted from [1]. B) **Mechanism of opposite tuning**. This diagram shows the activity of a group of opposite neurons lying on a ring. Orange line shows the background firing rate of opposite neurons. Red line shows how S2 input at *θ* inhibits neurons tuned to S1 inputs at *θ*. Under our model dynamics, this results in excitation of neurons tuned to S1 inputs at *θ* + 180° instead, as indicated by the green line. As such, opposite neurons are tuned oppositely to S1 and S2 inputs. C) **Model architecture**. Feedforward connections and recurrent connections are shown in the diagram, but divisive normalization is not shown. Arrow indicates excitatory connection, while a dot indicates inhibitory connection. Each group of neurons (S1, S2, C, O) are assumed to lie in a 1D ring formation, as shown in Fig. 1B. D) **A decentralized model of multisensory integration with opposite neurons**. Instead of having only one group of congruent and opposite neurons, each sensory modality has its own congruent and opposite neurons (C1, C2, O1, O2). Our current architecture in C) is equivalent to the boxed component, with S2 connection to C replaced by C2 connection to C1.

In order to circumvent this problem, we propose that instead of receiving excitatory inputs from oppositely tuned neurons, each opposite neuron receives an excitatory input and an inhibitory input from similarly tuned neurons. Suppose each opposite neuron is tuned to inputs at some angle *θ* from one of the sensory modalities S1, but receives inhibition from inputs at *θ* in the other sensory modality S2. An S2 input at *θ* then inhibits opposite neurons that are tuned to S1 inputs at *θ*. Under our model dynamics, this results in excitation of opposite neurons that are tuned to S1 inputs at *θ* + 180° instead. As such, opposite neurons are oppositely tuned to S1 and S2 inputs. See Fig. 1B. This formulation of opposite neurons is desirable as we no longer need to learn opposite connections, which is previously argued to be implausible due to the natural statistics of visual and vestibular inputs. Only Hebbian and Anti-Hebbian learning mechanisms are needed.

In some models of multisensory integration which incorporate congruent and opposite neurons, network dynamics are shaped by recurrent connections and divisive normalization [5, 6]. Our work aims to learn opposite neurons within the framework of these existing multisensory integration models, as an attempt to extend current theories. Our firing rate model employs Hebbian learning rules, and we show that the learned neurons have tuning properties reported in experiments. In addition, we demonstrate that these neurons can learn to be topographically organized, an essential property assumed in the aforementioned models of multisensory integration.

## 2 Materials and Methods

### 2.1 Network Architecture

Our work extends the decentralized model of multisensory integration proposed by Zhang et al. [5, 6], in which integration happens not in a single multisensory area, but is a result of communication between multiple, multisensory areas. We consider a simplified circuit with only one multisensory area, which could represent MSTd or VIP. Congruent (C) and opposite (O) neurons in this multisensory area receives input from two unisensory areas S1 and S2. An example of such a unisensory area would be the middle temporal area (MT), which is known to be the main visual input to MSTd [16]. A schematic of the model is shown in Fig. 1C. Inputs from S1 and S2 converge onto and excite the congruent neurons. S1 excites opposite neurons, while congruent neurons inhibits opposite neurons. Recurrent excitation and divisive normalization are present among the congruent and opposite neurons.

Why is this architecture chosen? As discussed in section 1, we need opposite neurons to receive excitation from one input and inhibition from the other, so a possible model architecture would involve excitation of opposite neurons from S1 and inhibition from S2. However, by Dale’s law, it is unlikely for S2 neurons to have excitatory projections to congruent neurons and inhibitory projections to opposite neurons at the same time. Therefore, instead of receiving inhibition from S2 directly, opposite neurons receive inhibition from congruent neurons, such that overall S2 still has an indirect inhibitory effect on the opposite neurons. This alternative arrangement also has a functional benefit: it allows us to justify adding a delay to the inhibitory signal to opposite neurons, which we find to be necessary for properly learning opposite neurons. See discussion of Eq. 7 for details.

As mentioned, recurrent excitation and divisive normalization are also present among the congruent and opposite neurons. These dynamics are present in previous decentralized models of multisensory integration [5, 6]. Recurrent excitation and divisive normalization are crucial for learning a topographical organization, and are based off the idea of Kohonen maps [17]. We note that other methods for achieving a topographical organization may also be possible, such as using a “Mexican-hat” shaped connection where there is local recurrent excitation and broad lateral inhibition.

Our model is not a decentralized model of multisensory integration, where there are separate groups of congruent and opposite neurons for each sensory modality. However, we note that the architecture can still be easily adapted to merge with a decentralized model by replacing input from S2 by input from C2, the congruent neurons for sensory modality 2 (Fig. 1D).

### 2.2 Network Dynamics

The model assumes that each group of neurons (S1, S2, C, O) lie on a 1D ring, with each neuron’s position parameterized by *θ* ∈ [−*π, π*). The mean firing rate of neurons in the sensory input areas S1 and S2 is given by

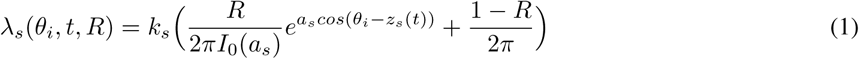

where *θ*_*i*_ is the position of neuron *i* on the 1D ring. The subscript *s* ∈ {1, 2} indicates whether the input is from S1 or S2. *I*_0_(*x*) is the modified Bessel function of the first kind with order 0. *k*_*s*_ is a scaling constant, while *R* ∈ [0, 1] is the *reliability* of the input. For example, for a visual self-motion input, *R* = 0.5 would correspond to 50% reliability/coherence of the random dot stimulus. *a*_*s*_ determines the width of the input, while *z*_*s*_(*t*) refers to the center of the input at time *t*.

This equation models the input as having the shape of a von-Mises distribution (first term of the equation) with a variable DC offset (second term of the equation). Von-Mises distribution can be thought of as an analogue of Gaussian distribution when the support is a circle. It is similar to the wrapped Gaussian distribution, which has been used to model the tuning of MSTd neurons [2]. Reliability controls the gain of the input but does not affect the input width. This is consistent with the observation that tuning bandwidth of MT neurons (which provides visual input to MSTd neurons) is roughly “coherence-invariant,” meaning it is invariant to changes in visual motion coherence [18]. The variable DC offset is set such that the total firing rate is independent of reliability, with 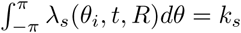. It is unclear whether MT neurons exhibit this property, but the requirement that total firing rate be relatively invariant to reliability is important to the functioning of our model. A comparison of the input in our model with MT neuron responses is shown in Figure 1 in the Supplementary Information.

The actual firing rate of neurons in S1 and S2 is then given by

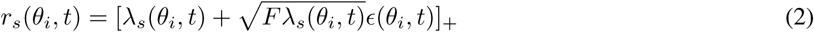

where *ϵ*(*θ*_*i*_, *t*) is Gaussian noise with *µ* = 0, *σ* = 1, [*x*]_+_ = max(*x*, 0), and *F* is the Fano Factor. We hereafter use 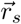 to represent the vector of input from S1 or S2.

Recurrent connections among congruent and opposite neurons are modeled by

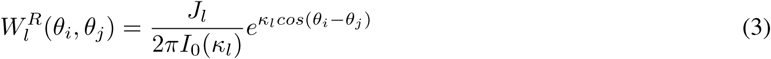

where *θ*_*i*_ and *θ*_*j*_ are positions of two different neurons *i* and *j* on the same ring. *l* ∈ {*c, o*} indicates whether the recurrent connection is among the ring of congruent or opposite neurons. We let 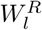 denote the matrix of connections. Note that *J*_*l*_ cannot be greater than a critical value *J*_*crit*_, or else the network can sustain a bump of activity indefinitely after removal of feedforward stimulus. The formula for *J*_*crit*_ has been derived by Zhang et al., and is given by

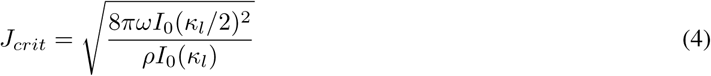

where *ρ* = *N/*2π [6].

We did not explicitly model divisive normalization using neurons. Instead, the effects of divisive normalization among the ring *l* of neurons are directly incorporated into the calculation of firing rate:

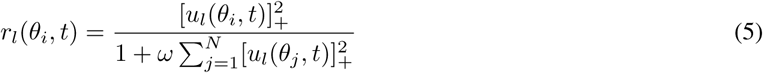

where *r*_*l*_(*θ*_*i*_) is the firing rate, *u*_*l*_(*θ*_*i*_) is the synaptic input, *N* is the number of neurons on the ring *l, ω* is a constant that controls the strength of normalization. The normalization operation described here was used by Carandini and Heeger to model divisive normalization observed in biological data [19]. It was also used in some previous studies of continuous attractor neural networks (CANN) [20, 21], as well as in the decentralized model of multisensory integration by Zhang et al from which our model follows [5, 6]. Experiments has also supported the presence of divisive normalization in multisensory integration areas [16]. We hypothesize that the operation could be carried out by a pool of inhibitory neurons. Again, we denote the vector of firing rates by 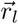 and the vector of synaptic inputs by 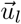.

Finally, letting *W*_*c*1_, *W*_*c*2_, *W*_*o*1_, *W*_*oc*_ be the feedforward connections from S1 to C, S2 to C, S1 to O, and C to O respectively, the dynamics of congruent neurons and opposite neurons are given by

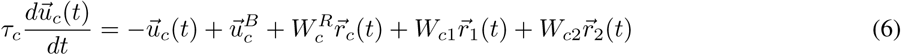

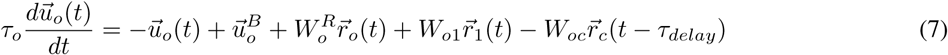

The first term on the right hand side is a decay term. The second term 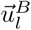, where *l* ∈ {*c, o*}, is a constant background input. The third term corresponds to input from recurrent connections, and 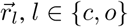 is given by Eq. 5. The fourth and fifth term correspond to feedforward inputs, with a negative sign in front of 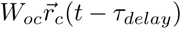 since the connection is inhibitory, as well as a delay *τ*_*delay*_ of the signal from congruent neurons to opposite neurons. This delay is essential for the learning of opposite neurons, for otherwise the excitatory and inhibitory input will cancel out increasingly throughout training, which in turn degrades the learning efficacy of opposite neurons. Adding the delay allows the excitatory input to be unaffected by the inhibitory input throughout the period of the delay, thus allowing opposite neurons to continue learning correctly. We note that this delay is also biologically plausible, since the inhibition, which goes through the congruent neurons first, is disynaptic, and would therefore be delayed in comparison to the monosynaptic excitatory input.

After each update, we rectify the weights with [*w*_*ij*_]_+_ to ensure all weights are non-negative.

### 2.3 Learning Rules

The network learns the feedforward excitatory and inhibitory weights via the same local, Hebbian learning rule, with

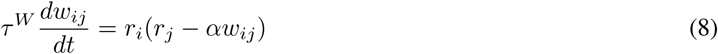

where *w*_*ij*_ denote an excitatory/inhibitory connection from neuron *j* to neuron *i*. We also enforce the constraint that all weights must be non-negative. Note that this is *not* Oja’s rule, where the second term inside the bracket would be *r*_*i*_*w*_*ij*_. Ursino et al. showed that for a simplified model without recurrent connections, this learning rule (with *α* = 1) for excitatory neurons allows the receptive field of the neuron to match its average input, which in turn allows maximum likelihood estimation in multisensory integration to be performed simply by reading out the position of the neuron with maximal firing rate [22]. The case of inhibitory neurons will be discussed further in Section 4.

### 2.4 Simulation Details

There were *N* = 180 neurons on each of the four rings of neurons, distributed uniformly over the stimulus space [−*π, π*) We set the synaptic input time constants to be *τ*_*c*_ = *τ*_*o*_ = *τ* = 0.01. Euler’s method was used with a step size of Δ*t* = 0.05*τ*, and the simulation was run for *T* = 120 = 240000Δ*t*. The learning rule time constant was *τ* ^*W*^ = 10 = 1000*τ*, and *α* = 15. For sensory inputs, *a*_1_ = *a*_2_ = 1.5, and *k*_1_ = *k*_2_ = 2π*I*_0_(3)*e*^−3^ ≈1.5. The position of input from S1, *z*_1_(*t*), was generated by first randomly permuting an evenly spaced sequence of inputs from −*π* to *π*, each lasting *τ*_*stim*_ = 10*τ*, then adding Gaussian noise with *µ* = 0 and *σ* = 2°. *z*_2_(*t*) was generated by adding Gaussian noise with the same *µ* and *σ* to *z*_1_(*t*). The Fano Factor *F* was set to 0.5. For the recurrent connections, *J*_1_ = *J*_2_ = 0.5*J*_*crit*_ (see Eq. 4), with *κ*_1_ = *κ*_2_ = 3. *ω* = 1.6*·*10^−2^ for divisive normalization.

All synaptic inputs and background inputs were initialized to 0. The feedforward weights were all initialized with the following method: Consider feedforward connections from a pool of input neurons indexed by *j* to a pool of target neurons indexed by *i*. For each *j*, we sample 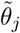 from a uniform distribution over the *N* target neurons without replacement (i.e. 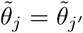 if and only if *j* = *j*′), as well as a multiplicative factor *A*_*j*_ from a log normal distribution with arithmetic mean of 3 and arithmetic variance of 1. Then the initial connections are given by

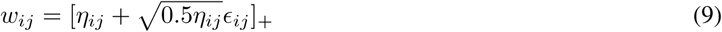

Where

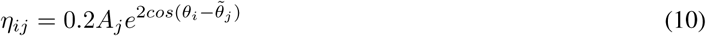

and *ϵ*_*ij*_ is i.i.d. Gaussian noise with *µ* = 0 and *σ* = 1. Intuitively, this models each input neuron as projecting to a random target location with variable connection strength and a spatial spread given by von Mises distribution. Figure 2 in the Supplementary Information shows the initialization of weights using this method.

**Figure 2:**
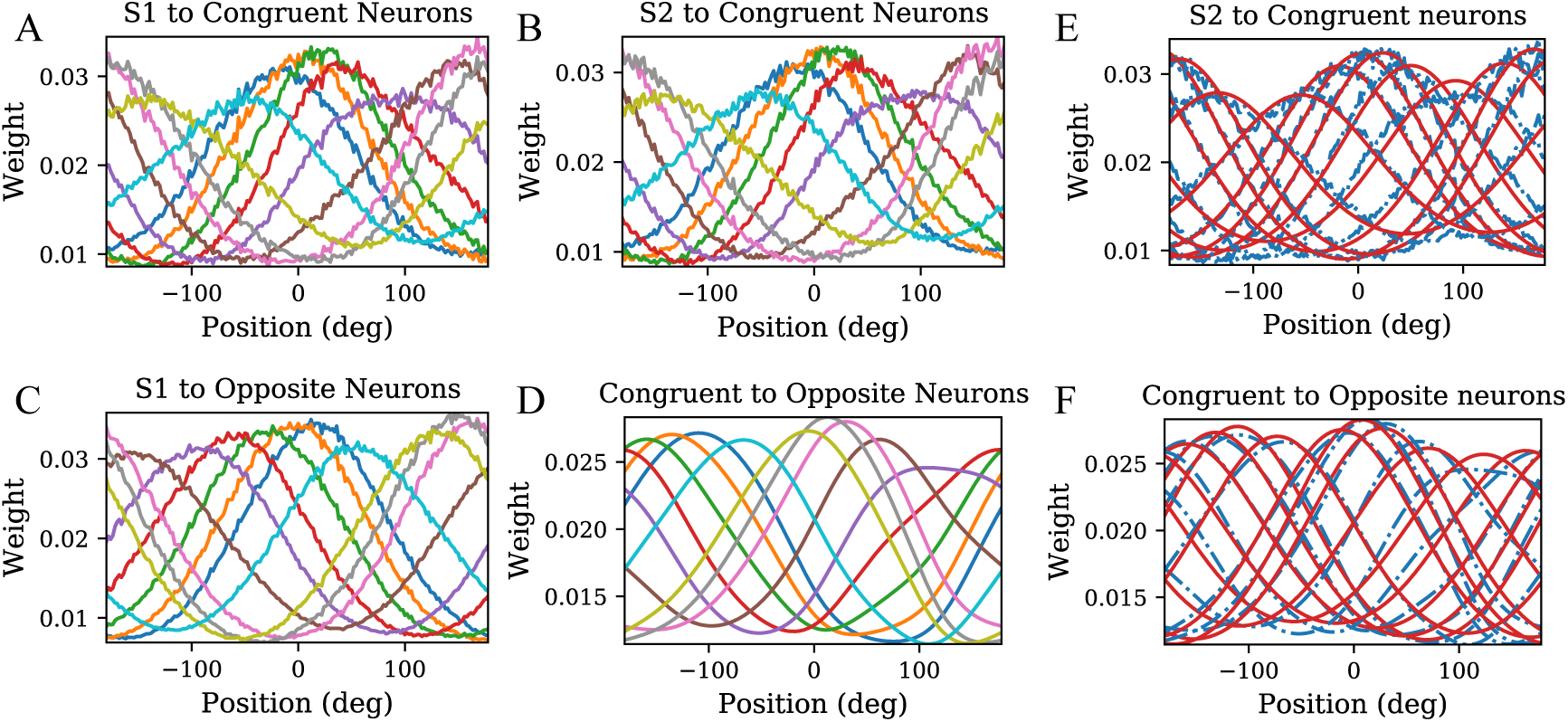
Feedforward weights have the shape of von-Mises distribution. A-D) Feedforward weights from S1/S2/congruent neurons to 10 congruent and opposite neurons. Each curve represents the feedforward weights to one congruent or opposite neuron. For D), note that although the connection strength shown here is positive, the connections are inhibitory because of the opposite sign in front of the 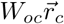 term in Eq. 7. E, F) Feedforward weights in B) and D) overlaid with best-fit von-Mises distribution. Blue curves are the feedforward weights, while red curves are the best-fit lines. The best fit curves are obtained by fitting them to von-Mises distributions scaled by some constant.

## 3 Results

Our simulation shows that our network is able to learn both congruent and opposite neurons, with tuning properties that agree with experimental data qualitatively.

### 3.1 Feedforward connections to congruent and opposite neurons

#### Feedforward weights have the shape of von-Mises distribution

We define the feedforward weights from S1/S2/congruent neurons to some neuron *i* as the weight vector 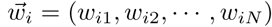, where *w*_*ij*_ is the weight of a connection from a S1/S2/congruent neuron *j* to neuron *i*. The shape of feedforward weights 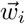 regarded as a function over its indices *j* is found to be approximately proportional to a von-Mises distribution. This is shown in Fig. 2. The assumption of Gaussian or von Mises shaped feedforward weights is usually assumed in multisensory integration models, and we show that our model can learn feedforward weights that have the same shape [5, 6, 23, 24, 22].

#### Topographic organization of congruent and opposite neurons

The congruent and opposite neurons we learned in our model are topographically organized. To demonstrate this, we plotted the feedforward weight matrices of congruent and opposite neurons in Fig. 3. The bright ridges that have slopes of either +1 or −1 show that the congruent and opposite neurons are organized in a clockwise or anti-clockwise manner with respect to the input space.

**Figure 3:**
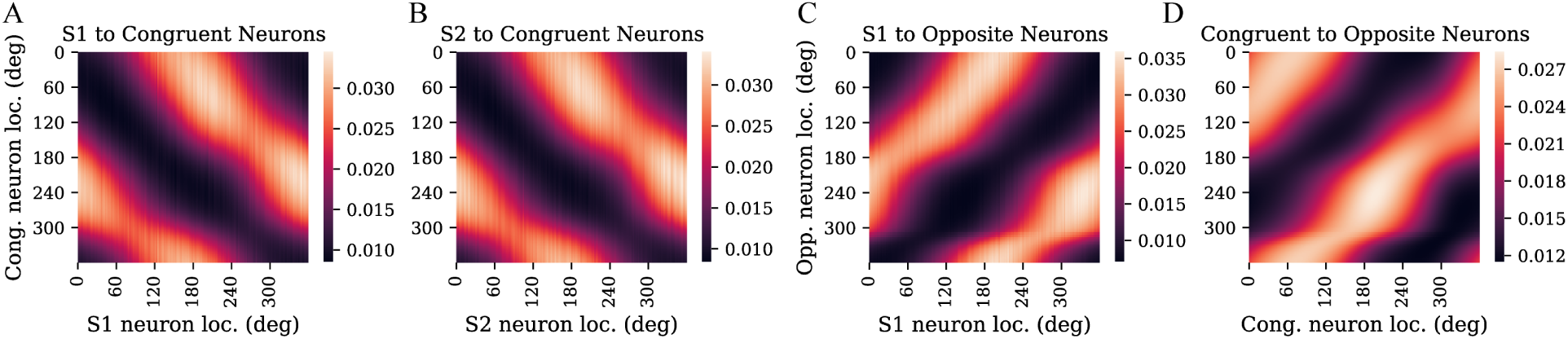
Topographic organization of congruent and opposite neurons. A-D) Weight matrices of feedforward connections to congruent and opposite neurons. The color represents the strength of the connection. In all cases, there is a bright ridge that has a slope of either +1 or −1, showing that the congruent and opposite neurons are organized in a clockwise or anti-clockwise manner with respect to the input space. A way to think about it is that as we move from 0° to 360° in the input space, we also complete a full revolution in the ring of congruent/opposite neurons.

### 3.2 Tuning curves of Congruent and Opposite Neurons

#### Tuning curves of congruent and opposite neurons to unimodal and bimodal stimuli

We mimicked psychophysical experiments and obtained the tuning curves of congruent and opposite neurons by varying the input stimulus location and recording the response of the neurons. We applied both unimodal and bimodal stimuli and compare the resulting tuning curves. Here, a unimodal S1 stimulus means that the mean firing rate at S1 follows Eq. 1 with *R* = 1, while the mean firing rate at S2 follows the same equation but with *R* = 0. Note that a unimodal S1 stimulus does not mean there is no input at S2, only that the input at S2 is constant. This is consistent with the observation that in MT neurons (which would correspond to S1 or S2 in our model) appear to have a non-zero background input [25, 18]. Moreover, we note that for our model to work, a certain kind of homeostasis must be maintained: the total input from S1 and S2 has to remain relatively constant. This necessitates the use of a constant background input at S2 when we assume a unimodal S1 stimulus. Further details are provided under Eq. 1. A bimodal stimulus means that both S1 and S2 neurons have the same mean firing rate with *z*_1_(*t*) = *z*_2_(*t*) (Eq. 1). The results are shown in Fig. 4A and B. From the figure, it can be seen that the congruent neuron has the same tuning to unimodal S1 and S2 stimuli. When bimodal stimuli are presented, the tuning of congruent neurons has a similar shape. Its response is stronger yet subadditive. For opposite neurons, tuning to unimodal S1 and S2 stimuli are separated by approximately 180°. When bimodal stimuli are presented, the response is flattened and highly sub-additive. The subadditivity of congruent and opposite neuron responses agree with experimental observations of MSTd neurons in macaques [2]. A qualitative comparison can be made with the experimental data shown in Fig. 4C-F, reproduced from [3].

**Figure 4:**
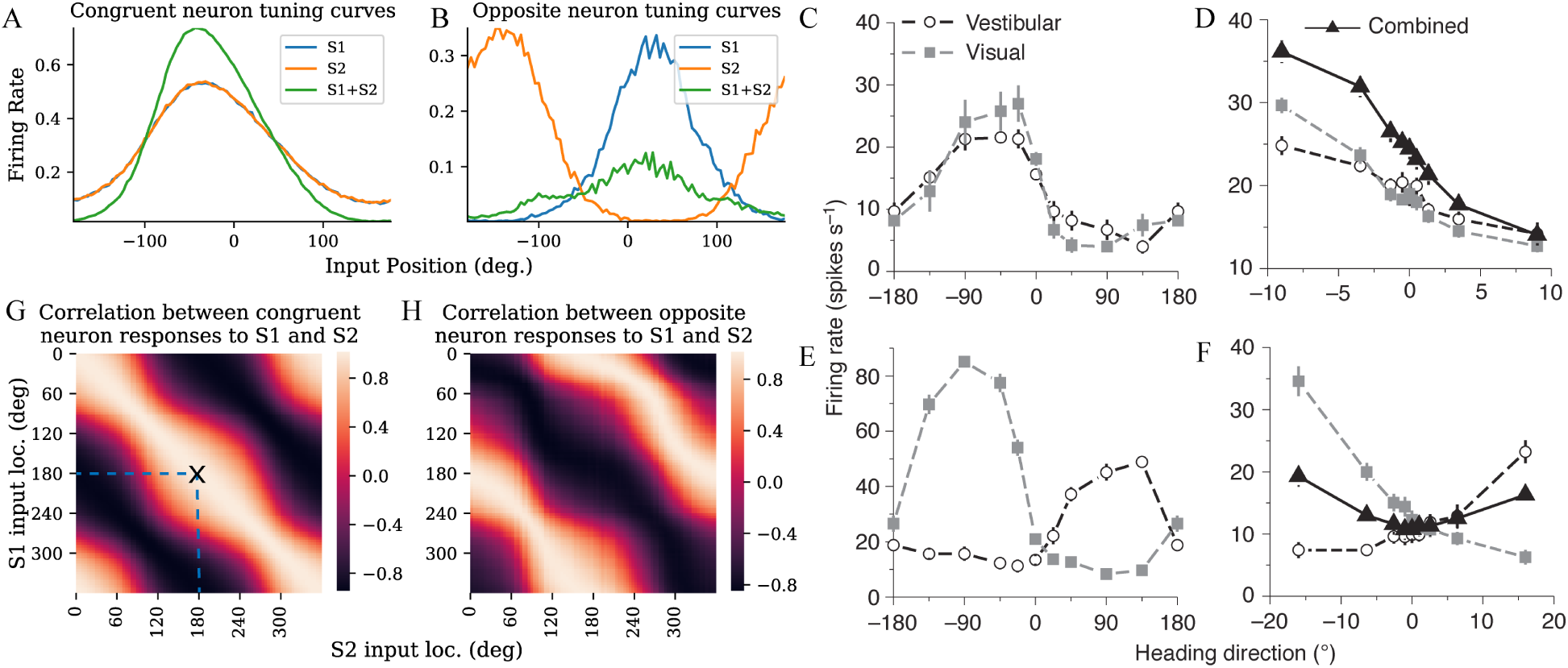
Tuning curves of congruent and opposite neurons to unimodal and bimodal stimuli. A) Tuning curve of a congruent neuron when a) only S1 stimulus is presented, b) only S2 stimulus is presented, and c) S1 and S2 stimulus are presented at the same location. B) Similar, but for opposite neuron. C-F) Experimental data of tuning curves of MSTd neurons to vestibular and visual stimulus, reproduced from [3]. C and D show the tuning of a congruent neuron, with D also showing the tuning of the neuron to combined vestibular and visual stimulus from 10° to −10°. E and F show the tuning of an opposite neuron. We see qualitative agreement between the tuning curves obtained from our simulation and tuning curves obtained from data. **Congruent and opposite neurons have congruent and opposite tunings**. G) Correlation between congruent neuron response to unimodal S1 stimulus and unimodal S2 stimulus. For example, the bright spot at X means that there is a strong correlation of congruent neuron response to S1 stimulus at 180° and S2 stimulus at 180°. We see a bright ridge along the main diagonal, meaning there is strong correlation of response to S1 and S2 inputs from the same location. H) Similar, but for opposite neurons. We see a bright ridge along the diagonal shifted by 180°, meaning there is strong correlation of response to S1 and S2 inputs from opposite locations.

#### Congruent and opposite neurons have congruent and opposite tunings

To show that congruent and opposite neurons have congruent and opposite tuning to S1 and S2 stimulus, we computed the correlation between congruent and opposite neuron response to unimodal S1 stimulus and unimodal S2 stimulus. The correlation between responses 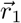 and 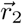 (both of which are vectors of firing rates) is measured by 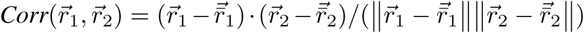, where 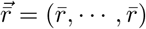, with 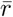 being the mean of 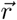. For congruent neurons, responses to unimodal S1 and unimodal S2 stimuli are most strongly correlated when the inputs are at the same location, while for opposite neurons, responses are most strongly correlated when the inputs are separated by 180°. The results are shown in Fig. 4G and H.

### 3.3 Effect of input reliability on tuning curve shape

#### “Contrast-invariant” tuning width of congruent and opposite neurons to unimodal stimuli

It is well-known that V1 neuron tuning is contrast-invariant [26, 27]. Similarly, a “coherence-invariant” tuning has also been observed in MT neurons, where the tuning width of MT neurons is more or less invariant to changes in the motion coherence of the random dot stimulus presented [18]. We demonstrate that the congruent and opposite neurons learned in our model also exhibit a similar property, namely that as we decrease the reliability of input from S1 or S2, the height of response from congruent and opposite neurons decrease, while maintaining the same tuning width. Reliability of a stimulus is encoded by the parameter *R* ∈[0, 1] in Eq. 1. For example, a random dot stimulus with 100% motion coherence will have a reliability of *R* = 1, while a random dot stimulus that is completely incoherent has *R* = 0. The results are shown in Fig. 5A.

**Figure 5:**
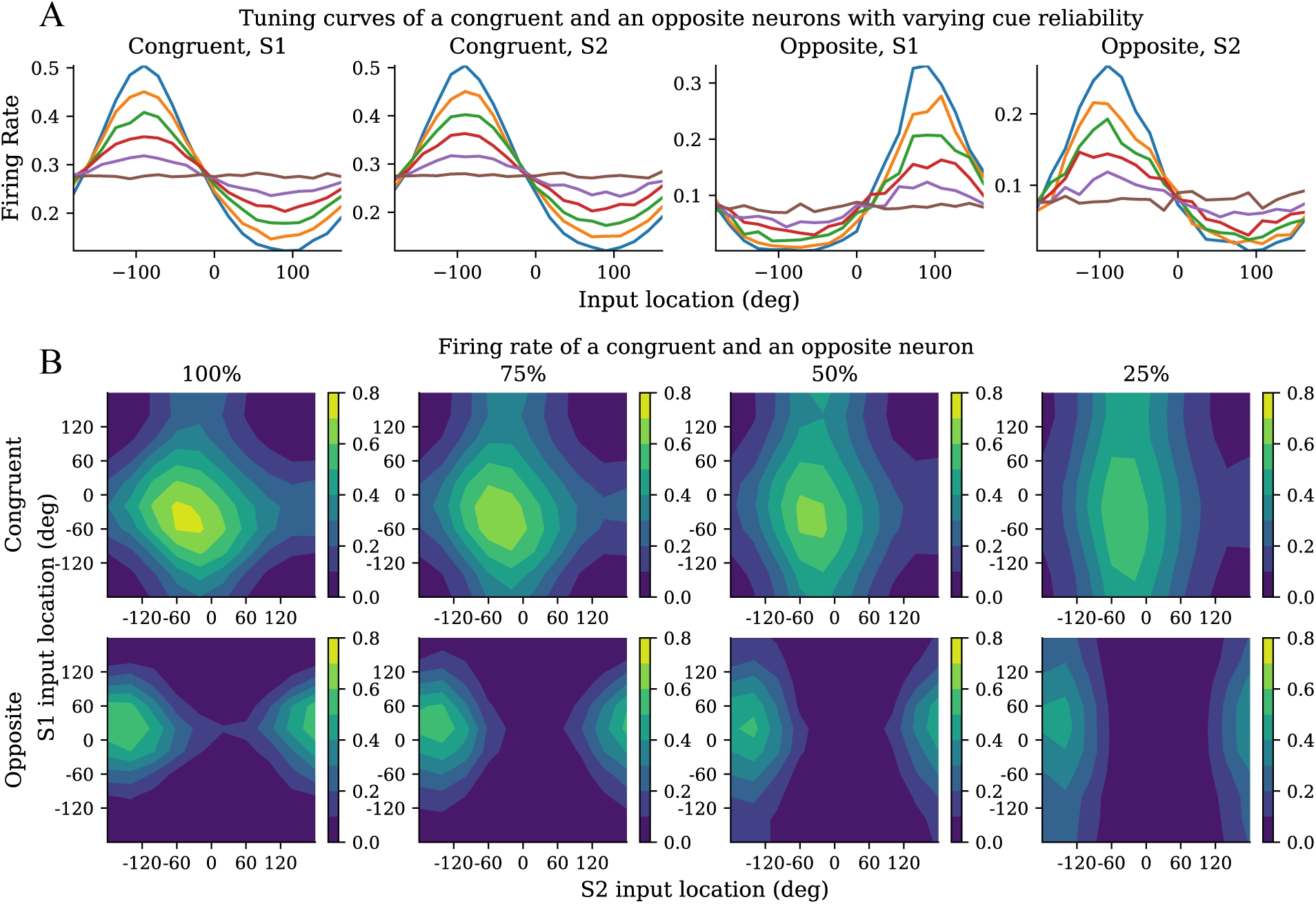
**A) “Contrast-invariant” tuning width of congruent and opposite neurons to unimodal stimuli.** Change in tuning curve of a congruent neuron and an opposite neuron to unimodal stimulus as input reliability is decreased. It can be seen that tuning width remains the same. This effect is due to recurrent excitation and divisive normalization in the circuit. Color labelling of reliability: blue - 100%, orange - 80%, green - 60%, red - 40%, purple - 20%, brown-0%. **B) Dependence of tuning on relative reliability of bimodal stimuli**. Change in tuning of a congruent and an opposite neuron to bimodal stimuli as S1 input reliability is decreased. As can be seen, decreasing S1 reliability shifts the tuning towards unimodal S2. Percentage indicates reliability of S1 input. Contour colors indicate firing rate.

#### Dependence of tuning on relative reliability of bimodal stimuli

Moreover, we show how the tuning of congruent and opposite neurons to both bimodal stimuli change as we decrease the reliability of one stimulus (Fig. 5B). Physiological experiments have shown that as the reliability of one stimulus decrease, the neuron should be increasingly tuned to the other, more reliable stimulus [2]. This effect is also observed in our model.

## 4 Discussion

### Summary and relation to other works

In this work, we used a biologically realistic rate-based model to learn opposite neurons that exhibit experimentally observed tuning properties to bimodal stimuli and are topographically organized. Our learned neurons display contrast-invariant tuning, a widely observed tuning property of V1 neurons [26, 27], and their response to varying reliability of input stimulus agree with experimental observations qualitatively. Our model architecture is compatible with some existing decentralized models of multisensory integration, and therefore our work also provides a basis for learning such models in general.

Some studies of ventriloquism have explored the learning of the equivalent of congruent neurons in a similar decentralized model [23, 24, 22], but they assume *a priori* topographic organization of the multisensory neurons before learning. Here, no such assumption is made, and topographic organization naturally emerges via a Kohonen map-like mechanism [17]. They also did not explore the learning of opposite neurons, which is the key contribution of our study.

### Anti-Hebbian learning rule

Anti-Hebbian learning, or learning of the inhibitory connections, in our model differs from some other rate-based models in which anti-Hebbian learning is involved [28, 29, 30]. Instead of assigning a different learning rule to inhibitory neurons, our inhibitory neurons follow the same Hebbian learning rule as the excitatory neurons. We speculate that such a simple learning rule worked for us because of the delay we introduced to the inhibitory signal from congruent to opposite neurons, which is biologically realistic because of our model architecture.

In fact, if such a delay is removed, opposite neurons cannot be learned well. We also tried using the anti-Hebbian learning rule introduced by Földiák, which takes the form Δ*w*_*ij*_ = −*α*(*r*_*i*_*r*_*j*_ −*p*^2^), for some fixed constant *p*. While opposite neurons can still be learned, the shape of the receptive fields can no longer be well-approximated by Gaussian or von-Mises distributions, an assumption in some decentralized models of multisensory integration [5, 6, 23, 24, 22].

### Causal Inference with opposite neurons

We are motivated by the theoretical observation that opposite neurons could provide a key step in causal inference by computing the Bayes factor [6, 15]. This is important because it is still unknown how the brain knows when to integrate or segregate multisensory cue information. However, the theoretical derivation assumes that opposite neurons simply sum up opposite inputs linearly, with no recurrent connections or divisive normalization among the opposite neurons. In contrast, our model is highly nonlinear [15]. Consequently, the theoretical derivations do not directly apply to the opposite neurons learned in our model. In future works, we aim to extend the theory to incorporate considerations of non-linearity in the circuit. Moreover, a possible future extension of this model would include a decision-making circuit that would determine when to integrate or segregation cue information based on the output of opposite neurons.

## 5 Conclusion

We have demonstrated in this paper that our model can learn opposite neurons that generally agree with experimental observations. Our congruent and opposite neurons also learn a topographic organization via a Kohonen map-like mechanism. In addition, our model can be easily integrated with some existing multisensory integration models, paving the way towards a complete circuit for performing multisensory integration that can optimally decide whether to combine or segregate cue information.

## Acknowledgments

This work was supported by NIH-funded Undergraduate Program in Neural Computation at the Center for the Neural Basis of Cognition.

