## Supplementary Information for "Emergence of opposite neurons in a firing-rate model of multisensory integration"

A PREPRINT

**Ho Yin Chau**
Department of Physics
University of California, Berkeley
Berkeley, CA 94720


**Wen-Hao Zhang**
Department of Mathematics
University of Pittsburgh
Pittsburgh, PA 15261


**Tai Sing Lee**
Center for the Neural Basis of Cognition
Carnegie Mellon University
Pittsburgh, PA 15289


October 21, 2019

### 1 Input at varying reliability

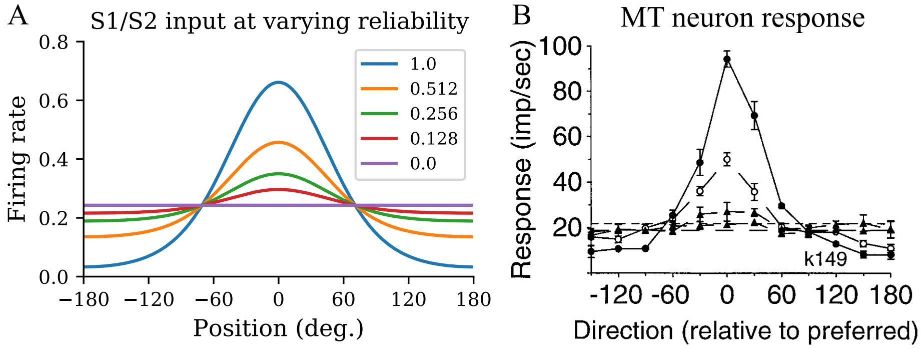


Figure 1: **Comparison of S1/S2 input at varying reliability with MT neuron responses.** A) S1/S2 input centered at $0°$ at varying reliability $R$, with value of $R$ indicated in the legend. B) Responses of one MT neuron to random dot stimuli at varying coherence levels. The coherence levels are $100\%$, $51.2\%$, $25.6\%$, and $12.8\%$, with $100\%$ being the curve with the strongest response. Background response of neuron, that is the activity of the same neuron when no stimulus is applied, is indicated by the flat dashed line. Figure is reproduced from [**?**]. Our inputs at S1 and S2 agree with the response of this particular MT neuron qualitatively.

### 2 Weight Initialization

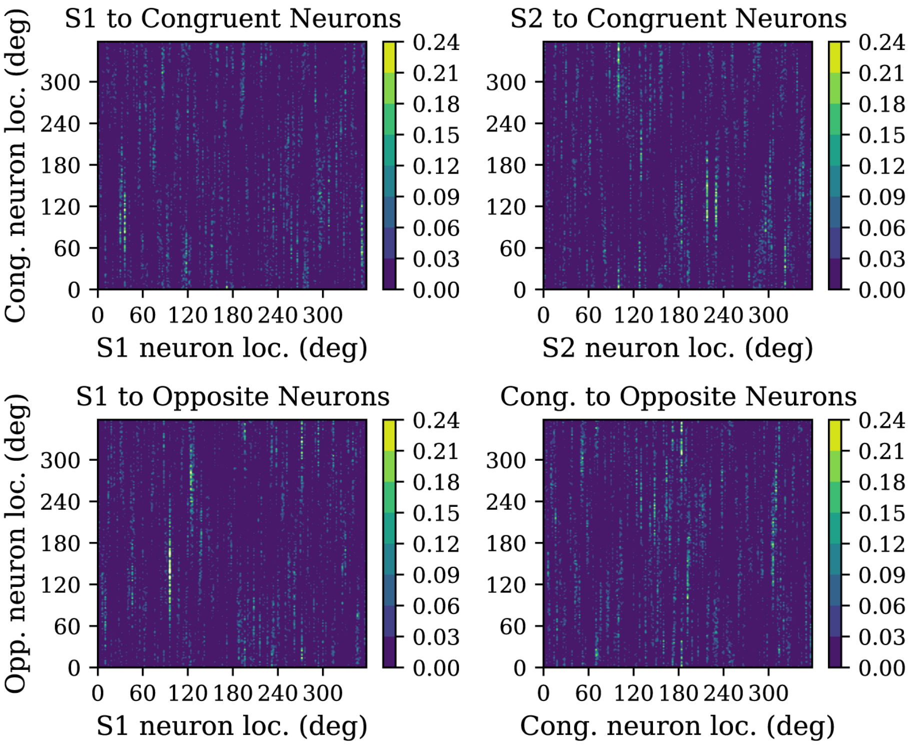


Figure 2: **Initialization of feedforward weights.** Each subplot shows the feedforward weight matrix at the beginning of training. As seen, there is no global structure in connectivity, and thus the topographical organization of weights at the end of training is learned. The initial weight distribution, however, is not homogeneous, with most weights being almost 0 along with localized regions with strong connectivity. The inhomogeneity is important for training as it causes some neurons to have much stronger activity than others, resulting in localized bumps of activity that encourages neurons next to each other to learn similar weights, which is needed for learning the correct topographical organization.
